# Genotype x Environment interaction and the evolution of sexual dimorphism: adult nutritional environment mediates selection and expression of sex-specific genetic variance in *D. melanogaster*

**DOI:** 10.1101/2023.08.15.553350

**Authors:** Stephen P. De Lisle

## Abstract

Sexual conflict plays a key role in the dynamics of adaptive evolution in sexually reproducing populations, and theory suggests an important role for variance in resource acquisition in generating or masking sexual conflict over fitness and life history traits. Here, I used a quantitative genetic genotype x environment experiment in *Drosophila melanogaster*, to test the theoretical prediction that variance in resource acquisition mediates variation in sex-specific component fitness. Holding larval conditions constant, I found that adult nutritional environments characterized by high protein content resulted in reduced survival of both sexes compared to an environment of lower protein content, and lower male reproductive success. Despite reduced mean fitness of both sexes in high protein environments, I found a sex*treatment interaction for the relationship between resource acquisition and fitness; estimates of the adaptive landscape indicate males were furthest from their optimum resource acquisition level in high protein environments, and females were furthest in low protein environments. Expression of genetic variance in resource acquisition and survival was highest for each sex in the environment it was best adapted to, although the treatment effects on expression of genetic variance eroded in the path from resource acquisition to total fitness. Cross-sex genetic correlations were strongly positive for resource acquisition, survival, and total fitness, and negative for mating success, although estimation error was high for all. These results demonstrate that environmental effects on resource acquisition can have predictable consequences for the expression of sex-specific genetic variance, but also that these effects of resource acquisition can erode through the life history.

## Introduction

The distribution of genetic variance for fitness and related traits plays an important role in governing the dynamics of adaptive evolution. In sexually-reproducing populations comprised of separate sexes, it is the bivariate distribution of male and female fitness that determines evolutionary rate (Lande 1980, Bonduriansky and Chenoweth 2009, Connallon and Hall 2016). When both sexes harbor substantial genetic variance in fitness and this variance is concordant – genotypes that have high fitness in one sex also tend to have high fitness when expressed in the other – then there is high potential for adaptive evolution. Alternatively, when genetic variance in fitness and related traits is sex-specific or sexually antagonistic, such that genotypes that have high fitness in one sex have low fitness when expressed in the other (Chippindale et al. 2001, Foerster et al. 2007), then this sexual conflict can impose a major constraint (Lande 1980) on the evolutionary process.

Simple life history models provide insight into the factors that mediate expression of genetic variance for fitness and its components. Assuming a life history can be defined by the acquisition of resources and subsequent allocation of resources to traits under selection, then variance in fitness can be decomposed into components related to variance in acquisition and variance in allocation strategy (van Noordwijk and de Jong 1986). Thus, key to the expression of genetic variance in fitness is the degree of variance (genetic or environmental) in resource acquisition. This framework applies directly to sexually reproducing populations (Zajitschek and Connallon 2017), where it is male and female genetic variance in resource acquisition that in part determines the degree of sex-specific genetic variance in fitness (Figure 1). There are also reasons to expect resource allocation strategies to be sex specific (Rowe and Houle 1996), which could further contribute to generating sex-specific variance in fitness.

**Figure 1.**
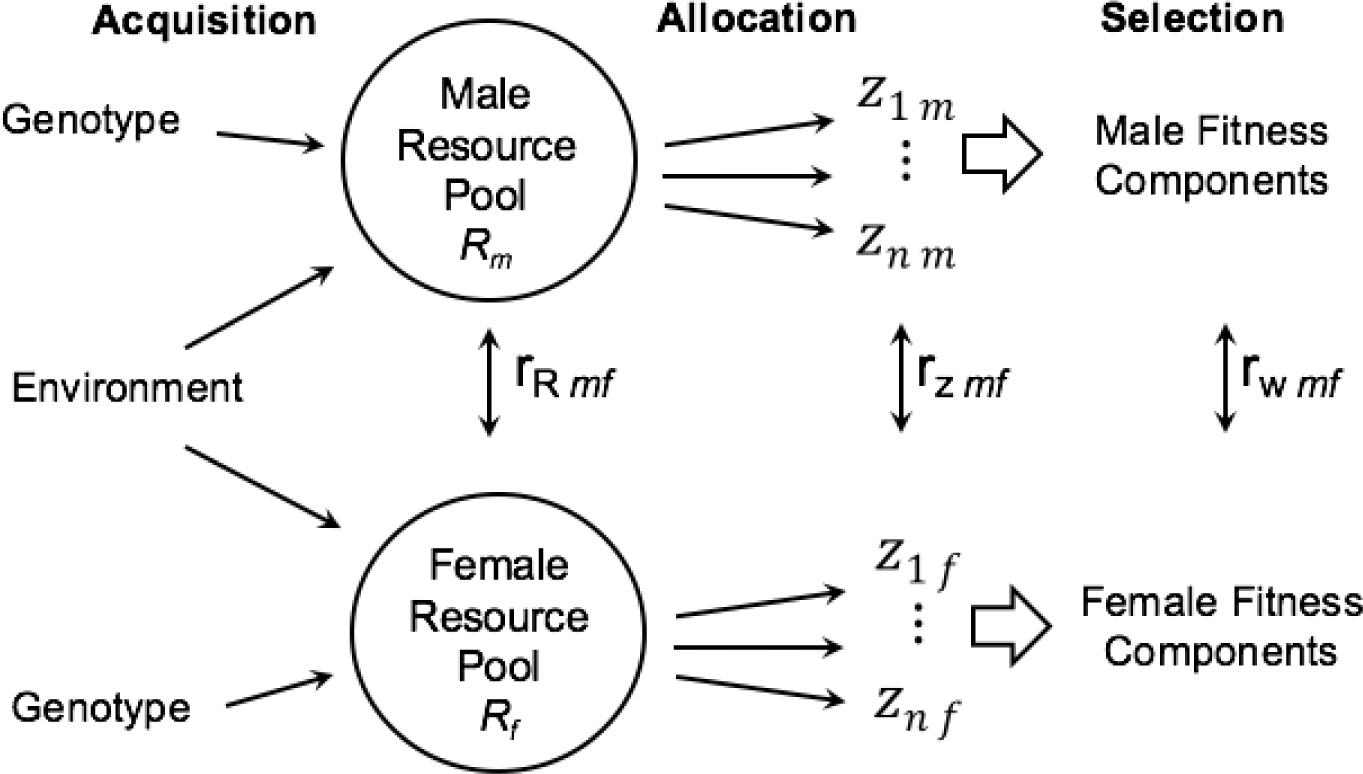
A graphical illustration of how (co)variance in resource acquisition and allocation determine cross-sex genetic correlations between fitness components (modified from Zajitscheck and Connallon 2017). Resources are acquired to determine the total resource pool (R) that an individual has available to allocate to a set of traits, *z_i-n_*, that are under selection. The genetic variance in resource acquisition and correlation between male and female resource acquisition has significant consequences to expression of genetic (co)variance in downstream traits, including *r*_z mf_ and most importantly, sex-specific fitness variance and the cross-sex genetic correlation for fitness *r*_w mf_. High values of *r*_R mf_ can swamp sexually-antagonistic selection on allocation to *z*, creating positive *r*_w mf_ and masking SA. Alternatively, the evolution of sex-differences in optimal resource acquisition could reduce *r*_R mf_, dramatically increasing sexual antagonism even if selection on allocation to *z* is not sexually antagonistic. Thus the model predicts that evolution of resource acquisition in males and females plays a central role in generating and resolving sexual conflict.

Although there are clear theoretical expectations for how sexual conflict for fitness is generated and impacts the dynamics of adaptive evolution, the empirical implications for actual organisms are less certain. Estimates of sex-specific genetic variance for fitness vary widely across populations and environmental conditions, even in the same study system (Barker et al. 2010, Gosden and Chenoweth 2014, Punzalan et al. 2014). Experiments in laboratory organisms a have shown that expression of genetic variance for fitness and component fitness can depend on the degree of adaptation to a given environment (Long et al. 2012), with the expectation that most genetic variance is concordant in situations of maladaptation (Connallon and Hall 2016). However, it is unclear how male and female nutritional environment may mediate variation in the expression of genetic variance. Importantly, many past laboratory manipulations of nutritional environment focus on resource acquisition prior to the onset of the expression of sexual dimorphism (e.g., larval food resources, Bonduriansky and Rowe 2005, Bonduriansky 2007, Punzalan et al. 2014, Holman and Jacomb 2017), making it difficult to assess the potential role that sex-specific resource acquisition may play in mediating sexual conflict.

Here I present results of a half-sib genotype x environment (GxE) quantitative genetic experiment in *Drosophila melanogaster* designed to understand three key questions. First, how does expression of genetic variance for fitness components change across adult nutritional environments which may differentially affect male and female fitness? Second, can environmental changes in the degree of sex-specific fitness variance be explained by shifts in genetic variance for resource acquisition, as predicted by life history theory? And third, how do these estimates of genetic variance relate to the adaptive landscape for resource acquisition in different nutritional environments? *Drosophila* flies are ideal for such a test; past work has demonstrated the importance of adult nutrition for both male and female fitness (Raubenheimer and Simpson 1997, Reddiex et al. 2013, Camus et al. 2017), as well variable degrees of sexual dimorphism in diet preference (Lee and Kim 2013, Davies et al. 2018, De Lisle 2023). My results illustrate how environmental shifts in expression of genetic variance for resource acquisition and component fitness can mediate the environmental dependence of sexual conflict.

## Materials and Methods

### Fly rearing and experimental design

Flies were obtained from a large stock population (LHm background; Rice et al. 2005, Lund-Hansen et al. 2020) maintained under standard conditions (25C, 12:12 light and 60% relative humidity) and density controlled each generation. A half-sib pedigree was set up by haphazardly selecting 60 newly-eclosed (virgin) male flies and placing them in a standard fly vial with 3 newly-enclosed females for 48 hours. Following this period of mating each female was then transferred to a new individual vial for egg laying for three days, after which females were removed from the vials. Offspring from these matings where then, upon eclosion, transferred to individual vials containing a single 5 μL microcapillary tube containing one of two liquid diets; a high-yeast diet containing 1:0.7 yeast:sucrose or a high sucrose diet containing 1:7 yeast:sucrose at a constant concentration of 0.1g/mL. These diet treatments correspond to protein:carbohydrate ratios of approximately 1:2 and 1:16 respectively, representing ends of a continuum of diet variation in *D. melanogaster* (Lee et al. 2008). Vials contained no regular fly food, although did contain approx. 5 mL of agar solution lacking nutrition to prevent desiccation. Four offspring from each genetic family (sire x dam combination; N = 135 that produced enough eclosed offspring for assay) were used; one male and one female in each respective liquid diet treatment. New microcapillary tubes were added to each vial after 48 and 96 hr. In order to control for effects of evaporation on measures of liquid food consumption, control vials with agar solution but no flies were also set up to obtain a baseline evaporation rate for each liquid food media. After 5 days of exposure to these diet treatments, all living flies were removed, remaining fluid in microcapillary tubes measured with digital calipers, and flies placed in new vials for competitive mating assays. These competition vials contained standard fly food and two additional flies, a male and a female virgin fly from the same genetic background as the stock population but homozygous for the *bw-* brown eye mutation (Chippindale et al. 2001). Following 24 hr in these in competition vials, all flies were removed and resulting offspring allowed to develop. The number of these offspring with *wt* eyes provided a measure of reproductive fitness.

### Statistical Analysis

To infer main affects of treatment on component fitness, I used two separate generalized linear models with sex, treatment, and their interaction as main effects. I assumed binomial error for analysis of survival, and Poisson error for offspring number for those individuals that survived. Analysis of the log odds of *wt*/*bw* offspring, instead of number of *wt*, produced similar conclusions, but this measure ignores fecundity variation and so is a less than ideal measure of fitness. To infer main effects on resource acquisition, I used a linear model (Gaussian error) with diet treatment, sex, and their interaction as fixed effects. This model contained fluid consumed, measured as the average of the fluid lost from each microcapillary tube a fly received, corrected for evaporation, as the response variable. Evaporation was corrected for by subtracting the average fluid lost from control fly vials containing microcapillaries with liquid food but lacking flies; separate corrections were performed for the two diet treatments to account for the differential evaporation of the different liquid diets.

I used a series of multi-response Bayesian mixed-effects models (Hadfield 2010) to estimate the two-sex **G** matrix separately for resource acquisition (gaussian), survival (threshold), reproductive success (Poisson), and total fitness (Poisson). Total fitness was calculated as the product of survival and reproductive success. For each model I used model comparison based on DIC of full (separate **G** for each nutritional environment) vs reduced (common **G**) models test the hypothesis that this matrix differs across adult nutritional environments. For survival, a binary trait, I fit a threshold model to estimate variance components but a categorical model for model comparison, as DIC is undefined for the threshold model. These models contained trait, treatment, and the trait*treatment interaction as fixed effects. Two G-side random effects were modelled; sire and dam*sire. To assess treatment effects on the **G** matrix, I fit two models differing in this G-side structure; one with a single sire level covariance matrix, and a second with separate matrices across low and high treatments. All G-side random effect covariance matrices were 2×2 matrices, with variances in male and female fitness components on the diagonal, and between-sex covariance of these fitness components on the off-diagonal. All models also included heterogenous residual variance; note that there is no defined covariance at this level as male and female fitness components were measured from different vials. Residual variance was fixed to unity for the threshold model of survival as this residual variance is undefined. For the cross-sex genetic correlation, *r*_mf_, I present pooled estimates from the reduced model for all traits as sampling error was high, making treatment-specific estimates difficult to interpret.

I used Poisson regression to infer phenotypic selection on resource acquisition, measured as the volume of liquid food media consumed by each fly. To assess statistical significance of treatment effects, I fit a single regression model with resource acquisition, treatment, and sex as fixed effects. This model included all two- and three-way interactions; interactions with resource acquisition indicate differences in phenotypic selection across treatments. This model, which already contained fixed-effect eight parameters, focused on directional selection only to avoid unnecessary complexity, although I estimated nonlinear selection gradients separately by treatment and sex (see below). I modelled absolute fitness as the response variable, noting that the log-linear Poisson model treats fitness on a log scale, analogous to relativizing fitness by the mean within each treatment*sex combination (De Lisle and Svensson 2017).

Following support for a treatment*sex differences in phenotypic selection, I estimated standardized directional and quadratic selection gradients separately for each treatment*sex combination. To estimate standardized selection gradients directly from the log-linear model, following Morrissey and Goudie (2022), I calculated linear and quadratic gradients as,

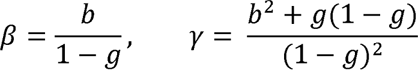

where *b* and *g* are the corresponding linear and (doubled; Stinchcombe et al. 2008) quadratic terms from a Poisson regression. I fit separate Poisson regressions to estimate *b* and *g* as the distribution of resource acquisition values was skewed (Lande and Arnold 1983). I mean-centered and variance standardized resource acquisition separately for each sex*treatment group. Standard errors for *β* and *γ* were obtained by resampling a multivariate normal distribution centered at the estimates of *b* and *g* with covariance equal to the information matrix from the fitted quadratic Poisson glm (Morrissey and Goudie 2022). Because all sex*treatment combinations showed a mixture of directional and stabilizing selection, it was appropriate to calculate the distance from the optimum phenotype, taken as *β*/-*γ* (Phillips and Arnold 1989), which gives the location of the optimum in units of phenotypic standard deviations from the mean. Full dataset and R script is available on Zenodo (https://doi.org/10.5281/zenodo.8248510).

## Results

The final dataset consisted of fitness measures from 534 individual flies (Figure 2A, B). Survival was lower for both sexes in the high protein environment (GLM z = 2.302, P = 0.021), and higher overall for males than for females in both environments (GLM z = 3.66, P = 0.000246; Figure 2C); there was no evidence of a sex*environment interaction for survival (GLM z = 0.012, P = 0.99). There were no main effects of treatment on the number of offspring produced in a competitive mating assay (GLM z = 1.608, P = 0.108), although there was a significant sex*treatment interaction (GLM z = 3.997, P = 6.41 x 10^-5^); males, suffered reduced mating success in high protein environments relative to low protein environments, although this effect was not observed in females (Figure 2D). Total fitness was thus lower in high protein environments than in low protein environments (Figure 2A, B). There was a sex*environment interaction for resource acquisition, measured as the volume of liquid media consumed in microliters (controlling for evaporation); females consumed more food in the low protein treatment than males, while both sexes showed similar, lower, levels of food intake in the high protein treatment (LM Sex* treatment t = −2.246, P = 0.0251; treatment t = 6.659, P = 6.9 x 10^-11^; Figure 3).

**Figure 2.**
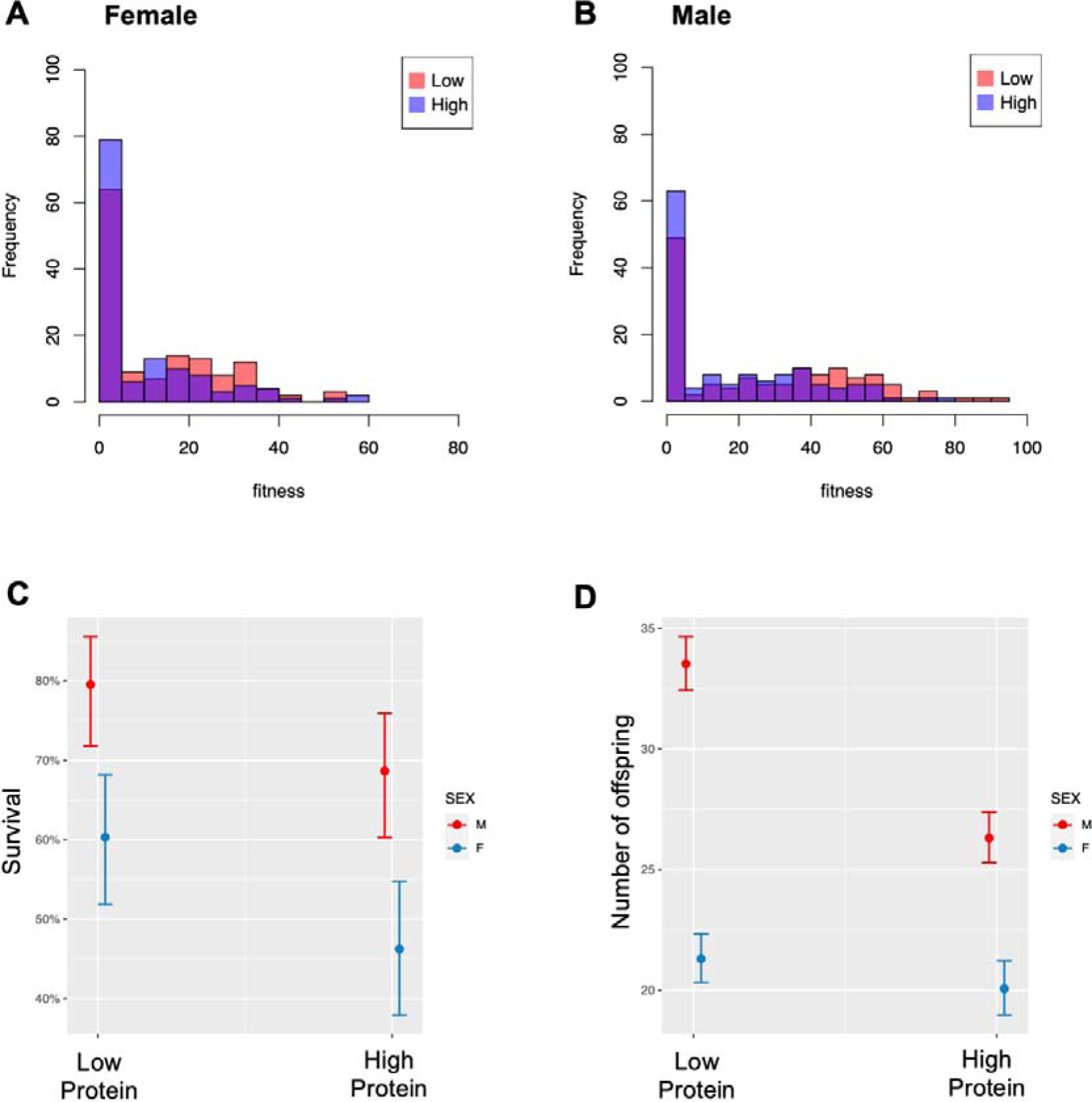
Total and component fitness under low and high protein adult nutritional environments. Panels A and B show distributions of total fitness for females and males in both nutritional environments. Survival was higher for males and in low protein environments (C). Fecundity was lower for males in high protein environments than in low protein environments, but did not differ for females (the sex*treatment interaction was significant). Panels C and D show marginal effects and 89% Cis from glms.

**Figure 3.**
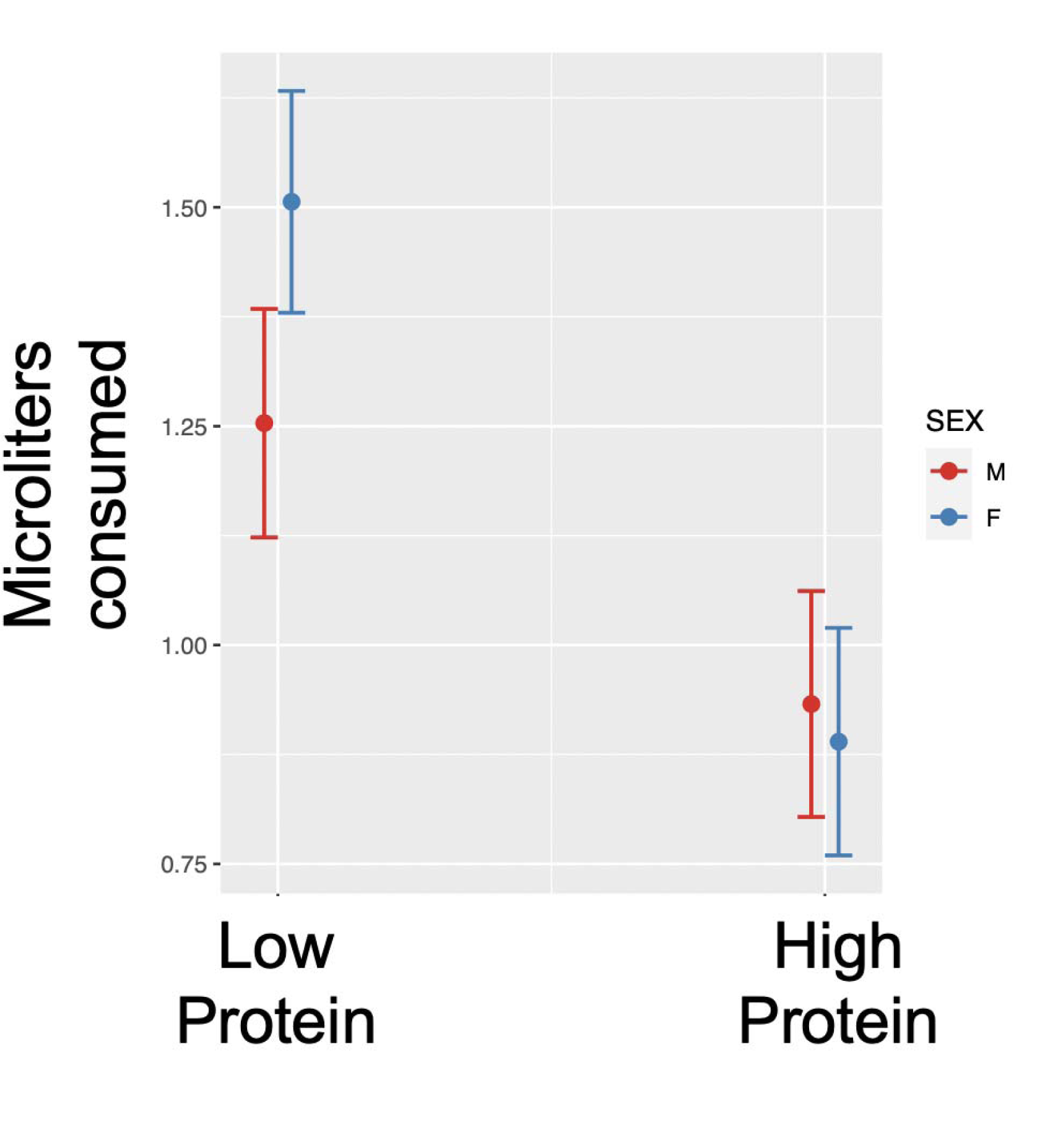
Food consumption in low and high protein nutritional environments. Both sexes consumed more liquid food media in low-protein conditions than in high protein conditions, although there was little evidence of a sex difference in consumption in high protein environments. Points are marginal effects and 89% CIs from a linear model.

Genetic variance, as captured in estimates of the univariate two-sex **G** matrix, differed substantially for resource acquisition and survival, but there was little evidence for treatment effects on **G** for reproductive success or total fitness (see Table 1). Female genetic variance in resource acquisition was higher in high protein environments than in low protein environments, while the opposite was true for males, which showed elevated levels of genetic variance in low protein environments (Figure 4). Although there was statistical support for environmental differences in G for survival, the effects were weaker (Figure 4). There was little statistical support for environmental differences in **G** for reproductive success or overall fitness (Table 1, Figure 4). Pooled (by treatment) estimates of the cross-sex genetic correlation, *r*_mf_, were strongly positive for resource acquisition, survival, and total fitness, and negative for mating success, although confidence limits for all estimates ranged from strongly negative to strongly positive (Table 2).

**Figure 4.**
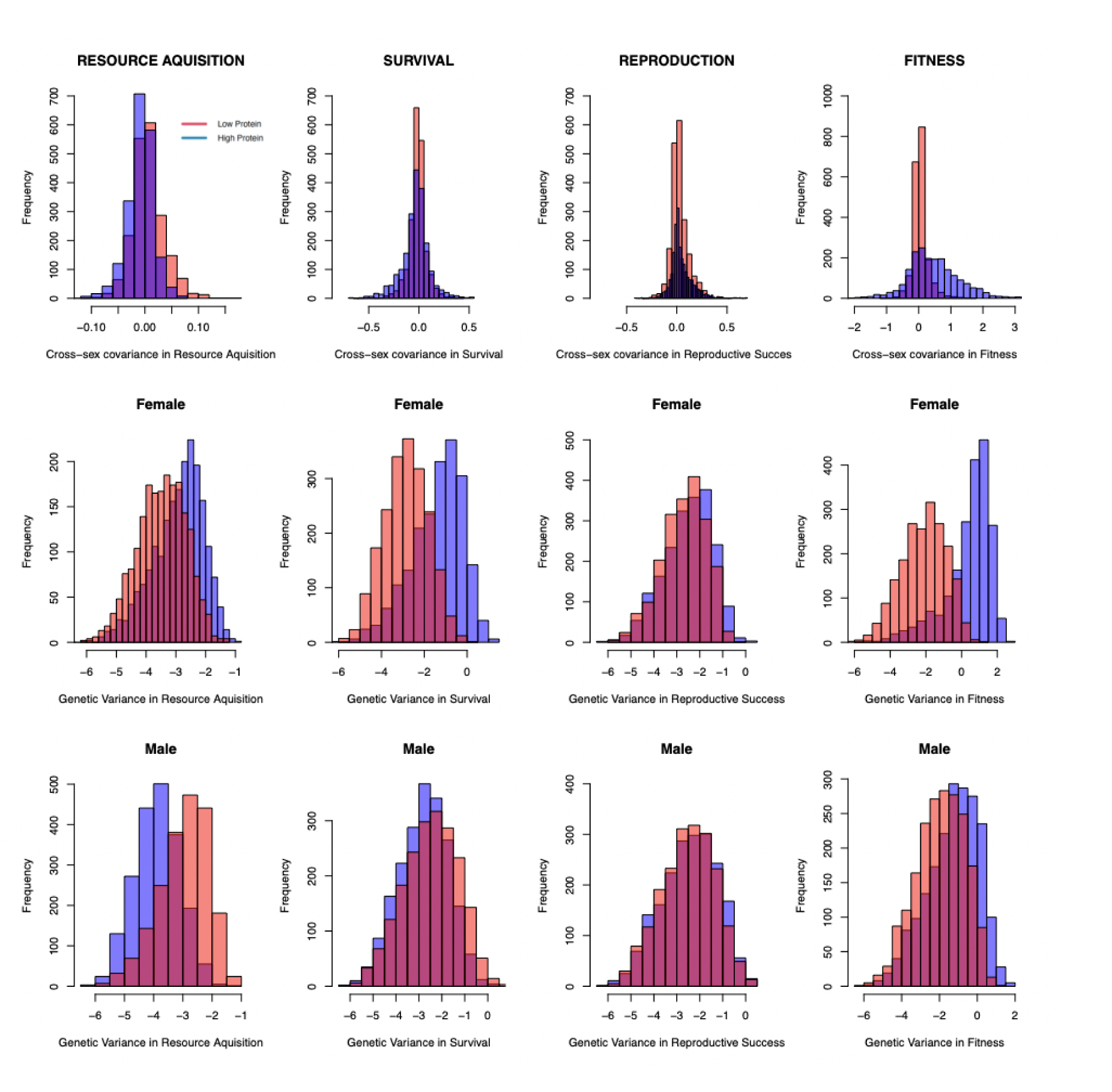
Posterior distributions of genetic variances for fitness components. Distributions show the either the raw posterior (for the cross-sex covariances in the top row) or the posterior on a log scale (for sex-specific variances) of the sire variance component for bivariate mixed models fit separately for each trait / fitness component. There was statistical support for treatment effects on the sire covariance matrix for resource acquisition and survival, but not for reproductive success or total fitness. Genetic correlations are presented in Table 2.

**Table 1.** Model comparison of multi-response mixed effects models to assess environmental differences in the two-sex G.

| Trait | DIC.Full | DIC.Reduced | $\Delta$ DIC |
| --- | --- | --- | --- |
| Resource Acquisition | 1196.844 | 1199.797 | -2.953236 |
| Survival | 641.0572 | 643.6579 | -2.600785 |
| Reproduction | 2136.301 | 2136.035 | 0.265237 |
| Fitness | 2239.584 | 2240.378 | -0.794426 |

**Table 2.** Estimates of cross-sex genetic correlations. Shown are the posterior mode and 95% highest posterior density interval from the reduced Bayesian mixed model for each trait.

| Trait | $r_{mf}$ | HPD interval |
| --- | --- | --- |
| Resource Aquisition | 0.667 | (-0.53, 0.91) |
| Survival | 0.908 | (-0.66, 0.99) |
| Reproduction | -0.897 | (-0.95, 0.85) |
| Fitness | 0.97 | (-0.90, 0.99) |

Selection on resource acquisition differed significantly by sex and environment (sex*environment*resource acquisition interaction, Poisson GLM z = −2.52, P = 0.011), although overall there was positive directional selection on selection on resource acquisition and negative quadratic curvature (stabilizing selection) of the fitness surface (Table 3), captured both in Poisson GLMs (Figure 5) and in estimates of standardized selection gradients calculated separately by sex and treatment (Table 4). Although each sex was displaced below its optimum in each environment (i.e. there was positive directional selection; Figure 5), males were closer to their optimum in the low protein environment whereas females were closer to their optimum resource acquisition level in the high protein environment (Figure 6). That is, there is evidence of sex-specific adaptation the different nutritional environments consistent with past work on nutritional requirements of insects, although it should be noted that sampling error for the location of the optimum was high despite statistical support for sex and treatment differences in selection.

**Figure 5.**
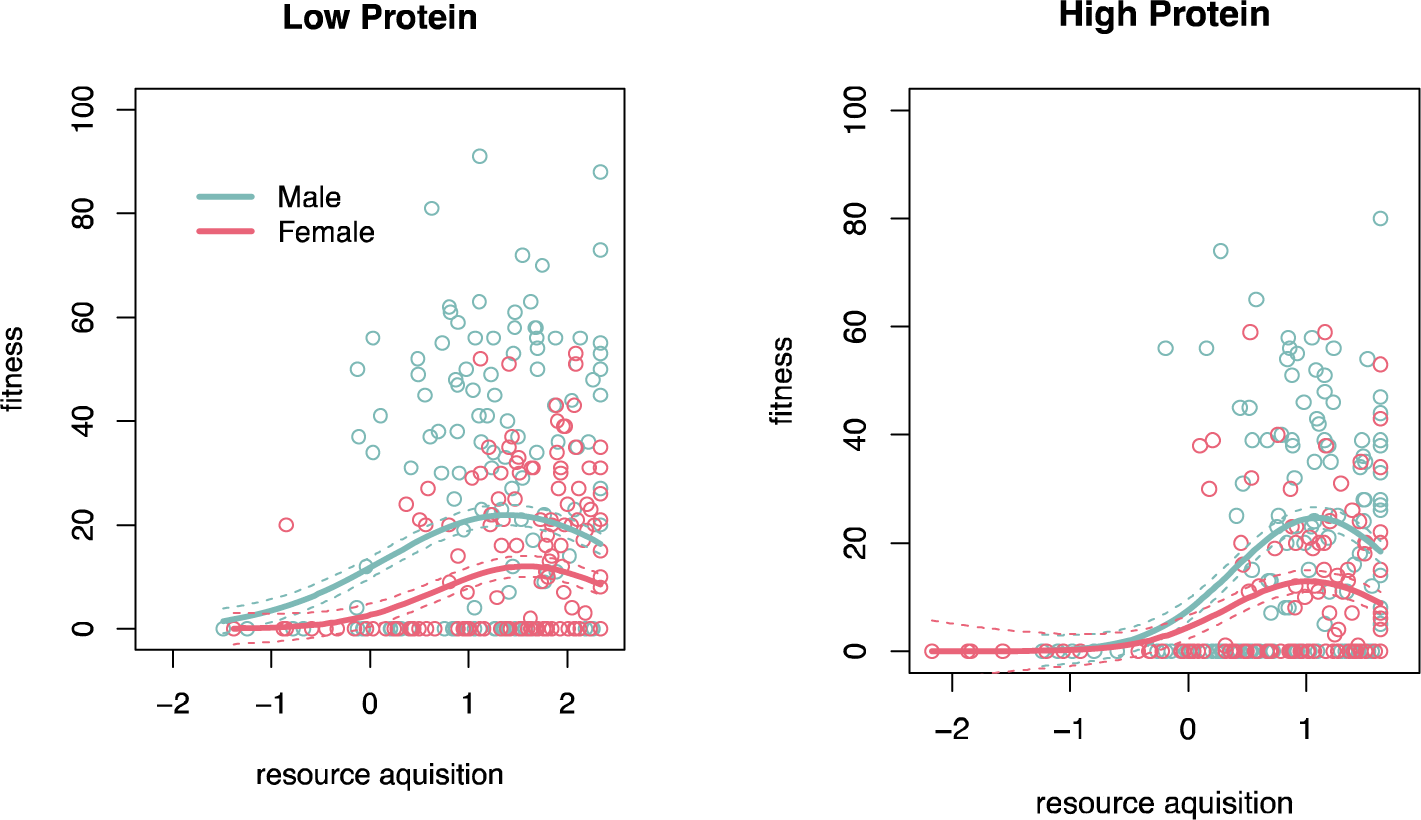
Individual fitness surfaces for male and female resource acquisition in low and high protein environments. Fitted lines show quadratic Poisson regressions fitted seperately for each sex*treatment combination. Similar conclusions on the shape of the fitness surfaces where obtained from cubic splines, suggesting the quadratic approximation is appropriate.

**Figure 6.**
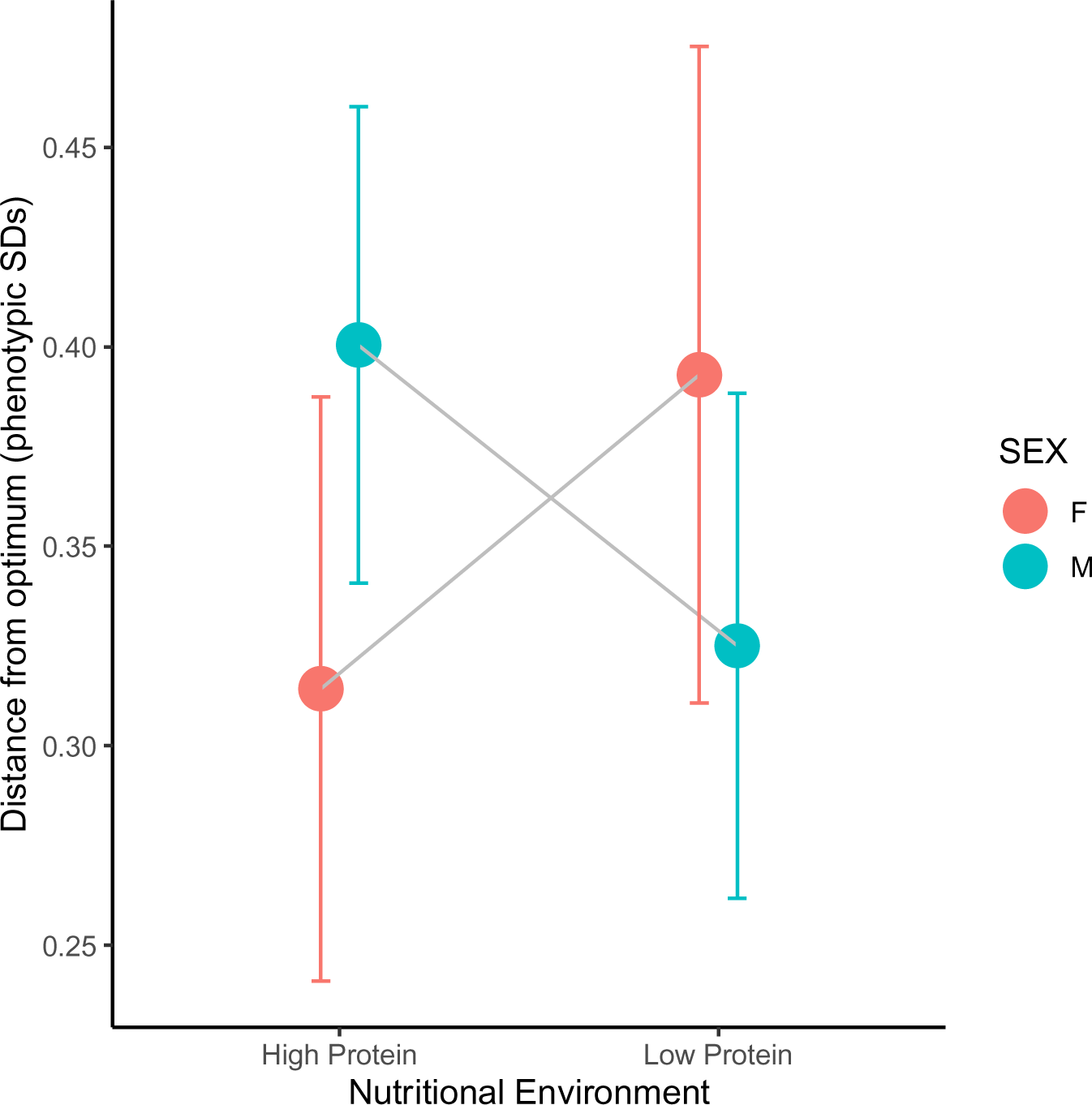
Distance from the optimum for each sex in each nutritional environment. Distance estimated as *β*/-*γ*, where variance standardized selection gradients were calculated from regression coefficients from Poisson regression (see text). Standard errors were calculated empirically by resampling the information matrix from the fitted Poisson regressions.

**Table 3.** Poisson GLM to assess treatment effects on sex-specific selection on resource acquisition.

| Effect | Estimate | SE | Z | P |
| --- | --- | --- | --- | --- |
| (Intercept) | 1.818759 | 0.057736 | 31.501 | <2x10<sup>-16</sup> |
| resource acquisition | 0.415489 | 0.045644 | 9.103 | <2x10<sup>-16</sup> |
| sex | 0.689204 | 0.071722 | 9.609 | <2x10<sup>-16</sup> |
| environment | 0.234847 | 0.084261 | 2.787 | 0.005318 |
| sex*resource acquisition | 0.009804 | 0.057808 | 0.17 | 0.865325 |
| treatment*resource acquisition | -0.088139 | 0.056758 | -1.553 | 0.120447 |
| sex*treatment | 0.339636 | 0.100289 | 3.387 | 0.000708 |
| sex*treatment*resource acquisition | -0.177709 | 0.070416 | -2.524 | 0.011612 |

**Table 4.** Standardized selection gradients for each sex*environment combination.

| **Sex** | **Environment** | **$\beta$** | **SE** | **$\gamma$** | **SE** |
| --- | --- | --- | --- | --- | --- |
| M | Low Protein | 0.090509 | 0.014874 | -0.278421 | 0.011012 |
| M | High Protein | 0.149633 | 0.015545 | -0.373672 | 0.017224 |
| F | Low Protein | 0.147805 | 0.020614 | -0.37612 | 0.024444 |
| F | High Protein | 0.143732 | 0.021335 | -0.45732 | 0.033975 |

## Discussion

Life history theory suggests that variance in fitness reflects a combination of variance in individual resource acquisition along with variance in how resources are allocated to traits under selection (van Noordwijk and de Jong 1986). For sexually reproducing organisms, patterns of sex-specific resource acquisition thus play a potentially important role in mediating the potential for sexual conflict for fitness (Zajitschek and Connallon 2017). Here I used a half-sib experiment in *Drosophila melanogaster* to understand how adult nutritional environment mediates the expression of resource acquisition variance, component fitness variance, and phenotypic selection in males and females. I found that each sex expresses greater genetic variance in resource acquisition rate in the nutritional environment they are best adapted to, as measured by distance from the phenotypic optimum resource acquisition level, and that this pattern of variance in resource acquisition manifests a similar pattern of genetic variance in survival. However, these treatment effects erode through the life history, with little evidence of treatment effects on genetic variance in mating success or total fitness. These data suggest that the nutritional environment can have important and predictable effects on the expression of genetic variance in fitness components.

I found that fitness was lower, and selection on resource acquisition stronger, for both sexes in the high protein nutritional environment. The observation of reduced survival for both sexes, and reduced competitive mating success for males, in high protein environments is generally consistent with past work on the nutritional fitness effects in flies and other insects (Maklakov et al. 2008, Reddiex et al. 2013, Garlapow et al. 2015, Jensen et al. 2015, Camus et al. 2017, Camus et al. 2018). In both environments, selection on male and female resource acquisition can be characterized as a combination of directional and stabilizing selection. This indicates that not only is resource acquisition under positive directional selection, as generally expected, but also that there exists an optimum level of resource acquisition within the vicinity of the phenotypic distribution. Despite consistent treatment differences in the strength of directional selection, male mean resource acquisition was closer to the optimum in the low protein nutritional environment, and further in high protein, while the opposite was true for females. This suggests that males are better adapted to low protein nutritional environments and females to high protein environments, which is consistent with past work in *D. melanogaster* (Reddiex et al. 2013, Camus et al. 2017) that used different experimental designs to assess selection on resource acquisition.

I found no evidence of a consistent male-bias in genetic variance, as has sometimes been reported for phenotypic traits (Wyman and Rowe 2014) and for fitness (Singh and Agrawal 2022). Estimates of cross-sex genetic correlations for all traits were high, either strongly positive (resource acquisition, survival, total fitness) or negative (reproductive success). This pattern suggests genetic variance is largely sexually concordant for both fitness and traits under natural selection, such as resource acquisition and survival, while the negative estimate of *r*_mf_ for competitive mating success suggests the presence of sexually antagonistic genetic variance for this fitness component consistent with past work (Chippindale et al. 2001). However, it is noteworthy that these genetic correlations are estimated with substantial error, with confidence regions widely overlapping zero. Nonetheless, the point estimates of *r*_mf_ are consistent with a scenario of elevated sexual conflict for mating success, with concordant variance for resource acquisition and survival.

Several caveats to the interpretation of these data are warranted. First, with fitness estimates from just over 500 flies from 60 sires, there was relatively limited power to infer treatment effects on genetic variance components, particularly cross-sex genetic correlations, which are notoriously challenging to estimate (Bonduriansky and Chenoweth 2009). Second, while fecundity was assessed in a competitive mating assay, thus capturing a measure of competitive reproductive success, this assay was performed only once, and so the contribution of late-life fecundity (Lee et al. 2008, Maklakov et al. 2008) was missed. Although this could be important for females, the assessment of reproductive fitness at approximately five days post-eclosure is consistent with the evolutionary history of LH_m_ flies. Third, feeding rates where determined by fluid loss from microcapillary tubes, a method subject to error arising from evaporation (De Lisle 2023), although evaporation was controlled for in calculations of resource acquisition.

Data and theory suggest that both selection and standing genetic variance may depended fundamentally on the location of a population relative to a phenotypic optimum. For example, when both sexes are maladapted, and thus far from their sex-specific optima, selection and genetic variance are expected to be sexually concordant (Connallon and Hall 2016), an expectation that seems to be supported by a range of empirical data from the lab and wild (Long et al. 2012, De Lisle et al. 2018). In this study, I was able to assess the effects of adaptation when both sexes are relatively close to their optima. Selection in this experiment was largely sexually-concordant, although subtle differences in the strength of directional selection, the curvature of quadratic selection, and differences male and female mean resource acquisition result in a situation where the sexes differ in the distance from their phenotypic optima in each environment, ultimately revealing that each sex is best adapted to a different nutritional environment. Estimates of genetic variance further reveal that variance in resource acquisition is highest for each sex in the environment they are best adapted to. Although it is unclear why this may be the case, it is noteworthy that the same pattern holds for genetic variance in survival, and for female mating success and total fitness. This finding of a concordance between variance in resource acquisition and component fitness is consistent with life history theory (van Noordwijk and de Jong 1986, Zajitschek and Connallon 2017), and the breakdown in the path to total fitness suggests other traits not associated with adult resource acquisition play a role in generating fitness variance. In *D. melanogaster*, this is to be expected, in part because most resource acquisition occurs at the larval stage.

This study also highlights the importance of inferring features of the adaptive landscape directly in making inferences about the degree of adaptation. Both sexes suffered reduced mean fitness in high protein environments, although this environmental affect on mean fitness does not entail a further displacement from the optimum resource acquisition level for females. Thus, although mean fitness was reduced, this reduction in mean fitness does not correspond to a large displacement of both sexes from the optimum resource acquisition level. This sort of discrepancy illustrates the utility of phenotypic selection analysis to infer features of the adaptive landscape (Lande and Arnold 1983, Arnold et al. 2001), even in laboratory studies of genetic variance.

As a proof of concept, these results partially support the theoretical expectation that sex-specific genetic variance in fitness components is related to variance in resource acquisition. More generally, the findings of environmental effects on sex-specific resource acquisition and sex-specific selection add to a growing body of work (Abruthnott et al. 2014, De Lisle 2019, 2023) demonstrating the role that ecological factors, such as diet and nutrition, can play in sex-specific adaptation.

## Acknowledgements

I thank K. Lund-Hansen for patient advice on *Drosophila*, and A. Singh, E. Svensson, and J. Abbott for sharing knowledge and resources. Petronella Romberg kindly assisted in setting up the experiment. This work was funded by grants from the Swedish Research Council (VR grant number 2019-03706 to SPD), Crafoord Foundation (20220602), Formas (grant 2021-01096), and the Royal Physiographical Society of Lund (Kungl Fysiografiska Sällskapet i Lund grants 42305, 41593) to SPD.

## References

Abruthnott, D., E. M. Dutton, A. F. Agrawal, and H. D. Rundle. 2014. The ecology of sexual conflict: ecologically dependent parallel evolution of male harm and female resistance in Drosophila melanogaster. Ecology Letters 17:221–228.

Arnold, S. J., M. E. Pfrender, and A. G. Jones. 2001. The adaptive landscape as a conceptual bridge between micro- and macroevolution. Genetica 112/113:9–32.

Barker, B. S., C. A. Phillips, and S. J. Arnold. 2010. A test of the conjecture that G-matrices are more stable than B-matrices. Evolution 64:2601–2613.

Bonduriansky, R. 2007. The evolution of condition-dependent sexual dimorphism. The American Naturalist 169:9–19.

Bonduriansky, R., and S. F. Chenoweth. 2009. Intralocus sexual conflict. Trends in Ecology & Evolution 24:280–288.

Bonduriansky, R., and L. Rowe. 2005. Sexual selection, genetic architecture, and the condition dependence of body size and shape in the sexually dimorphic fiy *Prochyliz xanthostoma* (Diptera: Piophilidae). Evolution:138–151.

Camus, M. F., K. Fowler, M. W. D. Piper, and M. Reuter. 2017. Sex and genotype effects on nutrient-dependent fitness landscapes in Drosophila melanogaster. Proceedings B 284:20172237.

Camus, M. F., C.-C. Huang, M. Reuter, and K. Fowler. 2018. Dietary choices are influenced by genotype, mating status, and sex in Drosophila melanogaster. Ecology and Evolution 8:5385–5393.

Chippindale, A. K., J. R. Gibson, and W. R. Rice. 2001. Negative genetic correlation for adult fitness between sexes reveals ontogenetic conflict in *Drosophila*. Proceedings of the National Academy of Sciences of the United States of America 98:1671–1675.

Connallon, T., and M. D. Hall. 2016. Genetic correlations and sex-specific adaptation in changing environments. Evolution 70:13.

Davies, L. R., M. F. Schou, T. N. Kristensen, and V. Loeschcke. 2018. Linking developmental diet to adult foraging choice in Drosophila melanogaster. Journal of Experimenal Biology 221:jeb175554.

De Lisle, S. P. 2019. Understanding the evolution of ecological sex differences: Integrating character displacement and the Darwin-Bateman paradigm. Evolution Letters 3:444–447.

De Lisle, S. P. 2023. Rapid evolution of sexual dimorphism driven by resource competition. Ecology Letters 26:124-131.

De Lisle, S. P., D. Goedert, A. M. Reedy, and E. I. Svensson. 2018. Climatic factors and species range positition predict sexually antagonistic selection across taxa. Philosophical Transactions of The Royal Society B. 373:20170415.

De Lisle, S. P., and E. I. Svensson. 2017. On the standardization of fitness and traits in comparative studies of phenotypic selection. Evolution 71:2313–2326.

Foerster, K., T. Coulson, B. C. Sheldon, J. M. Pemberton, T. H. Clutton-Brock, and L. E. B. Kruuk. 2007. Sexually antagonistic genetic variation for fitness in red deer. Nature 447:1107–1110.

Garlapow, M. E., W. Huang, M. T. Yarboro, K. R. Peterson, and T. F. C. Mackay. 2015. Quantitative genetics of food intake in Drosophila melanogaster. PloS OnE 10:e0138129.

Gosden, T. P., and S. F. Chenoweth. 2014. The evolutionary stability of cross-sex, cross-trait genetic covariances. Evolution 68:1687–1697.

Hadfield, J. D. 2010. MCMC methods for multi-response generalized linear mixed models: The MCMCglmm package. Journal of Statistical Software 33:1–22.

Holman, L., and F. Jacomb. 2017. The effects of stress and sex on selection, genetic covariance, and the evolutionary response. Journal of Evolutionary Biology 30:1898–1909.

Jensen, K., C. McClure, N. K. Priest, and J. Hunt. 2015. Sex-specific effects of protein and carbohyfrate intake on reproduction but not lifespan in Drosophila melanogaster. Aging Cell 14:605-615.

Lande, R. 1980. Sexual dimorphism, sexual selection, and adaptation in polygenic characters. Evolution 34:292-305.

Lande, R., and S. J. Arnold. 1983. The measurement of selection on correlated characters. Evolution 37:1210-1226.

Lee, K. P., and J.-S. Kim. 2013. Sexual dimorphism in nutrient intake and life span is mediated by mating in Drosophila melanogaster. Animal Behaviour 86:987–992.

Lee, K. P., S. J. Simpson, F. Clissold, R. Brooks, J. W. O. Ballard, P. W. Taylor, N. Soran, and D. Raubenheimer. 2008. Lifespan and reproduction in Drosophila: New insights from nutritional geometry. Proceedings of the National Academy of Sciences 105:2498–2503.

Long, T. A. F., A. F. Agrawal, and L. Rowe. 2012. The effect of sexual selection on offspring fitness depends on the nature of genetic variation. Current Biology 22:204–208.

Lund-Hansen, K. K., J. K. Abbott, and E. H. Morrow. 2020. Feminization of complex traits in Drosophila melanogaster via female-limited X chromosome evolution. Evolution 74:2703–2713.

Maklakov, A., S. J. Simpson, F. Zajitchek, M. D. Hall, J. Dessmann, F. Clissold, D. Raubenheimer, R. Bondurianksy, and R. C. Brooks. 2008. Sex-specific effects of nutrient intake on reproduciton and lifespan. Current Biology 18:1062–1066.

Morrissey, M. B., and I. B. J. Goudie. 2022. Analytical results for directional and quadratic selection gradients for log-linear models of fitness functions. Evolution 76:1378–1390.

Phillips, P. C., and S. J. Arnold. 1989. Visualizing multivariate selection. Evolution 43:1209–1222.

Punzalan, D., M. Delcourt, and H. D. Rundle. 2014. Comparing the intersex genetic correlation for fitness across novel environments in the fruit fly, Drosophila serrata. Heredity:143–149.

Raubenheimer, D., and S. J. Simpson. 1997. Integrative models of nutrient balancing: application to insects and vertebrates. Nutritional Research Reviews 10:151–179.

Reddiex, A. J., T. P. Gosden, R. Bonduriansky, and S. F. Chenoweth. 2013. Sex-specific fitness consequences of nutrient intake and the evolvability of diet preferences. The American Naturalist 182:91–102.

Rice, W. R., J. E. Linder, U. Friberg, T. A. Lew, E. H. Morrow, and A. D. Stewart. 2005. Inter-locus antagonistic coevolution as an engine of speciation: assessment with hemiclonal analysis. Proceedings of the National Academy of Sciences 102:6527–6534.

Rowe, L., and D. Houle. 1996. The lek paradox and the capture of genetic variance by condition dependent traits. Proceedings of the Royal Society of London Series B-Biological Sciences 263:1415–1421.

Singh, A., and A. F. Agrawal. 2022. Sex-specific variance in fitness and the efficacy of selection. The American Naturalist 199:587–602.

Stinchcombe, J. R., A. F. Agrawal, P. A. Hohenlohe, S. J. Arnold, and M. W. Blows. 2008. Estimating nonlinear selection gradients using quadratic regression coefficients: double or nothing? Evolution 62:2435-2440.

van Noordwijk, A. J., and G. de Jong. 1986. Acquisitiion and allocation of resources: Their influence of variation in life history tactics. The American Naturalist 128:137–142.

Wyman, M., and L. Rowe. 2014. Male bias in distributions of additive genetic, residual, and phenotypic variances of shared traits. The American Naturalist 184:326–337.

Zajitschek, F., and T. Connallon. 2017. Partitioning of resources: the evolutionary genetics of sexual conlflict over resource acquisition and allocation. Journal of Evolutionary Biology 30:826–838.

